# Effects of long-term closed environment on human saliva microbiota and salivary cytokines

**DOI:** 10.1101/2020.10.12.336750

**Authors:** Yinzhen Zhu, Zikai Hao, Yuming Fu, Jianlou Yang, Chen Dong, Hong Liu

## Abstract

Compared with the normal environment, the microbiota in controlled closed cabins such as space capsules, Lunar/Mars bases have changed. To ensure the health of crewmembers, it’s necessary to understand the effects of these changes on human symbiotic microorganisms and immunity. In this study, the experimental platform “Lunar Palace 1” with a similar closed and controlled environment was used to research the effects of changed microbial exposure on human saliva microbiota and salivary cytokines. This paper studied on four crewmembers who participated in the third phase of the “Lunar Palace 365” experiment, analyzing the dynamic changes of saliva microbiota and salivary cytokines, and further studying the correlation between salivary cytokines and highly abundant genera. According to our data, the crewmembers’ saliva microbiota and salivary cytokines fluctuated smoothly throughout the whole experiment. Although a part of microbes increased or decreased some times, they recovered quickly after leaving the controlled environment. The level of IL-6, IL-10 and TNF-α in crewmembers’ saliva decreased from normal environment to the controlled environment, showing reduced levels of oral inflammatory response in crewmembers. In addition, although there were significant individual differences in crewmembers’ saliva microbiota, sharing living space reduced the difference. Furthermore, the level of TNF-α showed a consistent positive correlation with the abundance of *Actinomyces* and *Rothia* in the controlled environment, indicating healthy individuals’ oral mucosal barrier may be sensitive to changes in saliva microbiota. According to the result, semi-sterile environments in controlled closed cabins didn’t cause persistent changes in human saliva microbiota and oral immunity. Besides, it provides a new idea for future research on the impact of the controlled environment on crewmembers health, and provides guidance for studying the effect of semi-sterile environments on human immunity based on saliva microbiota.

**Key points:**

1. Saliva microbes kept stable for individual but got convergent when sharing space;
2. The level of salivary cytokines reduced after entering the controlled environment;
3. There were complex correlations between salivary cytokines and saliva microbes;
4. The crewmembers adapt well to the controlled environment.

## Introduction

Long-term space missions and the prospect for future life in space promote research on the health of astronauts(White and Averner 2001). Studies have found that extreme pressure environments such as microgravity, cosmic radiation and semi-sterile environments have an effect on the human microbiome, the human immune system and the intricate balance between both, causing impaired immunity and increased susceptibility(Crucian et al. 2014, Gueguinou et al. 2009, Mermel 2013, Taylor 1993). The first two factors have been widely concerned, but the impact of closed isolation environment on symbiotic microorganisms and astronauts’ health is only noticed recently(Saei and Barzegari 2012).

Oral cavity is the gateway for pathogens and toxicant to invade human. Oral microbiota plays a vital role in maintaining oral and general health. Oral mucosal epithelial cells and dendritic cells can distinguish symbiotic and pathogenic microorganisms by pattern recognition receptors such as Toll-like receptors, and mediate immune inflammatory response to potential invading pathogens or immune tolerance to systemic microorganisms(Feller et al. 2013, Moutsopoulos and Konkel 2018). Disorders of oral microbiota not only cause various oral infectious diseases, but also relevant to digestive diseases, cardiovascular diseases, diabetes, rheumatoid arthritis, et.al.(He et al. 2015, Scannapieco 2013). Therefore, researching the chang of astronauts’ oral microbial community in space missions can provide some guidance for revealing oral and general health of astronauts.

Recently, studies find that long-term space flight can cause change in astronauts’ oral microbiota, causing oral and systemic inflammatory diseases. In Skylab missions, moderate increases were observed in the in-flight increments of dental plaque, calculus, and gingival inflammation, and increased counts of oral *Streptococcus*, *Neisseria*, *Lactobacilli*, and *Enteric bacilli* was detected after flight(Brown et al. 1976). Besides, other studies found that astronauts may suffer from conjunctivitis, upper respiratory tract infections, viral gastroenteritis, rhinitis and skin infections(Taylor and Sommer 2005). In the past, it was thought that the main factors causing it were microgravity and radiation, and the effects of semi-sterile environments in closed isolation cabins on human health was unclear.

Bioregenerative Life Support Systems (BLSS) can provide sustainable life support in a closed artificial ecosystem and is the future direction of space stations and planetary bases(Drake et al. 2010, Zheng et al. 2008). Lunar Palace 1 (LP1) is one of the most advanced BLSS, offering an unique confined environment. It is a good experimental model for studying the effects of space capsules on human systemic microorganisms and immune disorders. LP1 has the following characteristics: (1) It is a closed experimental system with no material exchange with the outside world, which is beneficial to maintain environment microorganisms stable in the system; (2) Weekly disinfection effectively prevents microbial reproduction and provides a semi-sterile environment for the crewmembers(Sun et al. 2016); (3) Crewmembers work according to a fixed schedule and maintain a good psychological situation(Hao et al. 2019). We suspect that long-term living in this semi-sterile environment will reduce cremembers’ exposure of environment microorganisms, which may cause salivary microbial change and oral inflammatory response reduce. When cremembers return to the normal environment, increased microbes may cause saliva microbiota alter and oral inflammatory response increase.

To demonstrate our hypothesis, we explored the effect of semi-sterile environment on cremembers’ symbiotic microbes and immune system in BLSS. We monitored saliva microbiota and salivary cytokines of crewmembers before entering LP1, during living in LP1, and after leaving PL1. This paper mainly studied the changes of crewmembers’ saliva microbiota and salivary cytokines as well as their relationship in BLSS, and discussed the impact of environmental microbial decreasing on it.

## Materials and methods

### Participants

The participants in this study were four Chinese crewmembers involved in the 3^rd^ Phase of the “Lunar Palace 365” project, including two males (Subject A and B) and two females (Subject C and D). All crewmembers had no history of smoking, oral disease, serious illness and chronic disease. Most of their general physical examination indices (performed at the 306th Hospital of PLA, Beijing, China) before and after 3^rd^ Phase were within normal ranges.

### Study design

The ‘‘Lunar Palace 365” project was a 370-day, multicrew, closed experiment carried out in a ground-based experimental BLSS platform named LP1. Located at the Institute of Environmental Biology and Life Support Technology, Beihang University, Beijing, China, LP1 was a highly closed ecosystem integrating efficient higher plant cultivation, animal protein production, urine nitrogen recycling, and bioconversion of solid waste; it had achieved an overall closure coefficient of 97% in terms of mass regeneration(Fu et al. 2016). Then we upgraded LP1 in 2016. Currently, it consists of a comprehensive cabin and two plant cabins with a total area of 160 m2 and a total volume of 500 m3. The comprehensive cabin includes four private bedrooms, a living room, a bathroom, and an insect culturing room (Fig 1(a)). The ‘‘Lunar Palace 365” project was divided into three phases, completed by two groups (group 1 and group 2) (Fig 1(b)). We explored the temporal dynamics of the saliva microbiota and salivary cytokines in the group 1 participating to the 3^rd^ Phase of the ‘‘Lunar Palace 365” project, including 1 month before entering LP1 and 1 month after returning to regular life. The experiment was designed as Fig 1 (c).

**Fig 1:**
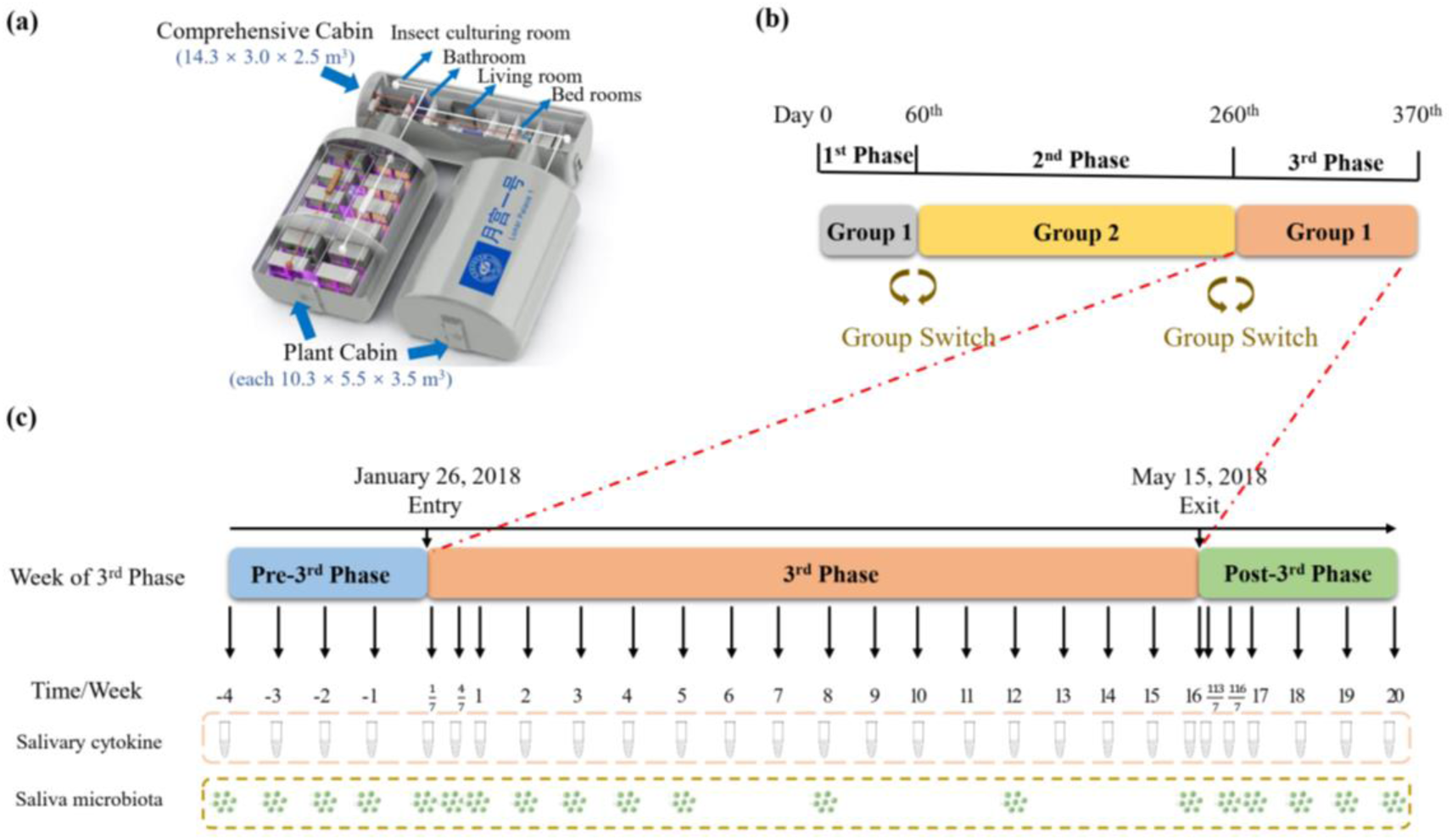
(a) Structure of “Lunar Palace 1”, (b) The mission of “Lunar Palace 365” project, (c) Experimental design: Tube:crewmembers’ salivary cytokines sampling; Eight green dots: crewmembers’ saliva microbiota sampling.

### Collection of whole saliva and determination of cytokine in saliva

All saliva samples were collected from 9:30 pm to 10:30 pm to minimize circadian variation in salivary composition. The crewmembers were not allowed to eat or drink anything except water and to perform oral hygiene activities including tooth brushing before collection. Saliva was collected using the spitting method for 10 min. Cellular debris in saliva samples were removed by centrifuging at 4000× g for 20 min at 4 ℃, and supernatants were aliquoted and stored at −80 ℃ until further analyses. Cytokines in saliva were quantified using Enzyme-linked Immune Sorbent Assay (IL-1β, IL-6, IL-10, TNF-α, IFN-γ, RGB&CHN).

### Collection of whole saliva and Illumina HiSeq sequencing

Saliva samples were collected at the time set in Fig 1 (c). Except for water, nothing should be eaten within 2 hours before sampling, and oral hygiene activities should not be carried out. Saliva samples were collected as follows: (1) Relax cheeks and gently rub for 30 seconds to produce saliva, (2) Spit saliva into a 5 ml cryotube and collect about 1 ml of saliva, (3) Add 2 ml saliva preservation solution to the cryotube and mix it by inversion 10-20 times, (4) Store the cryotube in −20 °C freezers for further testing.

Saliva samples were taken from the −20 ℃ refrigerator every three months and defrosted for 30 minutes. Total bacterial DNA were extracted from samples using the Power Soil DNA Isolation Kit (MO BIO Laboratories) according to the manufacturer’s protocol. The V3-V4 region of the bacterial 16S rRNA gene was amplified with the common primer pair (Forward primer, 5’-ACTCCTACGGGAGGCAGCA-3’; reverse primer, 5’-GGACTACHVGGGTWTCTAAT-3’) combined with adapter sequences and barcode sequences. PCR amplification was performed in a total volume of 50 μl, which contained 10 μl Bμffer, 0.2 μl Q5 High-Fidelity DNA Polymerase, 10 μl High GC Enhancer, 1 μl dNTP, 10μM of each primer, 50 ng genome DNA, and the remaining volume was supplemented with ddH2O. Thermal cycling conditions were as follows: an pre-denaturation at 95 °C for 5 min, followed by 15 cycles denaturation at 95 °C for 1 min, renaturation at 50 °C for 1 min and extension at 72 °C for 1 min, with a final extension at 72 °C for 7 min. The PCR products from the first step PCR were purified through VAHTSTM DNA Clean Beads. A second round PCR was then performed in a 40μl reaction which contained 20 μl 2×Phμsion HF MM, 8 μl ddH2O, 10μl of each primer and 10μl PCR products from the first step. Thermal cycling conditions were as follows: an pre-denaturation at 98 °C for 30s, followed by 10 cycles denaturation at 98 °C for 10s, renaturation at 65 °C for 30s and extension at 72 °C for 30s, with a final extension at 72 °C for 5 min. Next, all PCR products were quantified by Quant-iT™ dsDNA HS Reagent and pooled together according to the quality ratio 1:1. Finally, agarose gel electrophoresis was performed at 1.8%. High-throughput sequencing analysis of bacterial rRNA genes was performed on the purified, pooled sample using the Illumina Hiseq 2500 platform(2×250 paired ends) at Biomarker Technologies Corporation, Beijing, China.

The raw sequence was spliced and filtered using FLASH (version 1.2.11) (Tanja and Salzberg 2011) and Trimmomatic (version 0.33) (Bolger et al. 2014), successively, and then the chimera was removed using UCHIME (version 8.1) (Edgar et al. 2011)to obtain high quality Tags sequences. The quality checked 16S rRNA gene sequences were classified into operational taxonomic units (OTUs) within a 0.03 difference (equivalent to 97% similarity) using USEARCH (version10.0) (Robert 2013). Taxonomic classification at different taxonomic levels of these OTU sequences was done with RDP classifier (version 2.2, http://sourceforge.net/projects/rdpclassifier/)(Wang et al. 2007) against the Silva (Release128, http://www.arb-silva.de)(Quast et al. 2012) and UNITE (Release 7.2, http://unite.ut.ee/index.php)(Kõljalg et al. 2013) using a 80% confidence threshold. Construct a phylogenetic tree at the genus level based on PyNAST(version 1.2.2, http://biocore.github.io/pynast/) (Caporaso et al. 2010) and ClustalW2 http://www.ebi.ac.uk/Tools/msa/clustalw2/)(Larkin et al. 2007), and then calculate the distance matrix among samples.

Sequencing-based 16S rRNA surveys are usually normalized by converting OTU sequence counts into fractional abundances for each sample. However, this standard technique leads to what is known as compositional effects(Jonathan and Eric 2012), and may cause false relationships between OTUs, or between OTUs and cytokines. For the analysis of the saliva microbiota dynamics over the whole experiment, the normalization technique developed by David et al. (David et al. 2014) was used. Briefly, for each crewmember: (i) Time points were normalized in the standard manner so that the sum of all fractional OTU abundances at a given time point was 1; (2) Highly abundant OTUs, accounting for 90% of median time point reads, were selected; (3) each time point was normalized to a reference community that was computed for each sample based on other time points with a similar community structure. Specifically, reference OTU values were computed using a weighted median across time series, with time point weights set to be (1 − *j*)^2^ and *j* being the pairwise Jensen-Shannon Distance (JSD) score to the sample being normalized.

### Statistics

All statistical analyses were conducted with R and MATLAB. As crewmember’s saliva microbiota and salivary cytokines are susceptible to other biotic and abiotic units, theoretically a set of hypothetical stochastic differential equations could be used to express the influencing mechanisms of biotic and abiotic factors on the dynamic response of crewmembers’ saliva microbiota and salivary cytokines. If a well-designed BLSS could provide sufficient sustenance support for the crewmembers, they could be acclimate themselves to the closed environment. Under this assumption, the crewmembers’ saliva microbiota and salivary cytokines variations should be a stationary stochastic process. We adopted the autocorrelation function (ACF) (McMurry and Politis 2010, 2015) to testify the above assumption by evaluating whether the ACF would be only dependent on the time interval, rather than time. If they were stationary stochastic process, we used Mann-Kendall trend test (Libiseller and Grimvall 2002, McCuen 1995) to analyze whether there is a trend change from the normal environment into LP1. As for the differences among groups in saliva microbiota, principal component analysis (PCA) and multivariate analysis of variance (MANOVA) were performed using R packages (reshape 2, ggplot 2 and vegan) and MATLAB (the MathWorks Inc.), respectively. Further, we analyzed the alpha and beta diversity of saliva microbiota at different time points and different phases. Depending on the normality and variance homogeneity of the data, the One-Way ANOVA or Kruskal-Wallis rank sum test was used, and *P* values were corrected for multiple testing using the Bonferroni method. The relationships among saliva microbiota and salivary cytokines were analyzed with Spearman’s correlation tests using R (ver 3.5.3) packages (hmisc), then adjusted for multiple comparisons with the FDR correction method.

## Results

### Individual differences and stability of saliva microbiota

The saliva microbiota of the four crewmembers (subject A to D) were tracked over time during the third phase of the “Lunar Palace 365” experiment. A total of 70 saliva samples were collected. Each sample was measured by 16S rRNA gene Illumina HiSeq sequencing (V3-V4 region), obtaining a total of 3,888,658 pairs of high-quality sequence reads. After splicing and filtering, a total of 3,312,409 Clean tags were generated (minimum of all sample, 17,811; mean of all sample, 33,820). The dynamics of each crewmembers’ oral microbial communities were reconstructed over time according to the normalization strategy described by David et al.(David et al. 2014). The most abundant phyla were *Actinobacteria, Bacteroidetes, Firmicutes, Fusobacteria, Proteobacteria, Spirochaetes* and *SR1*, accumulating more than 97% in most samples (Fig 2(a)). We selected the OTUs that existed at more than half of the time points as core OTUs for dynamics analysis. We obtained 97, 104, 111, and 108 core OTUs in subject A, B, C and D, respectively (Supplementary Table S1). As shown in Figure 2, these dynamic changes revealed the differences among individuals were much larger than variation within individuals over time. We further visualized the individual and gender differences of saliva microbiome using principal component analysis (PCA) (Fig 3(a-b)). Interestingly, individual and gender differences were differentiated on the PCA score plots (Fig 3(a-b)). We further performed multivariate analysis of variance (MANOVA) to statistically compare individual and gender differences between different groups. Hierarchical clustering was generated based on mahalanobis distances as a result of MANOVA (Fig 3(c-d)). The MANOVA analysis showed that there were significant differences (*p* < 0.001) between each individual and gender.

**Fig 2:**
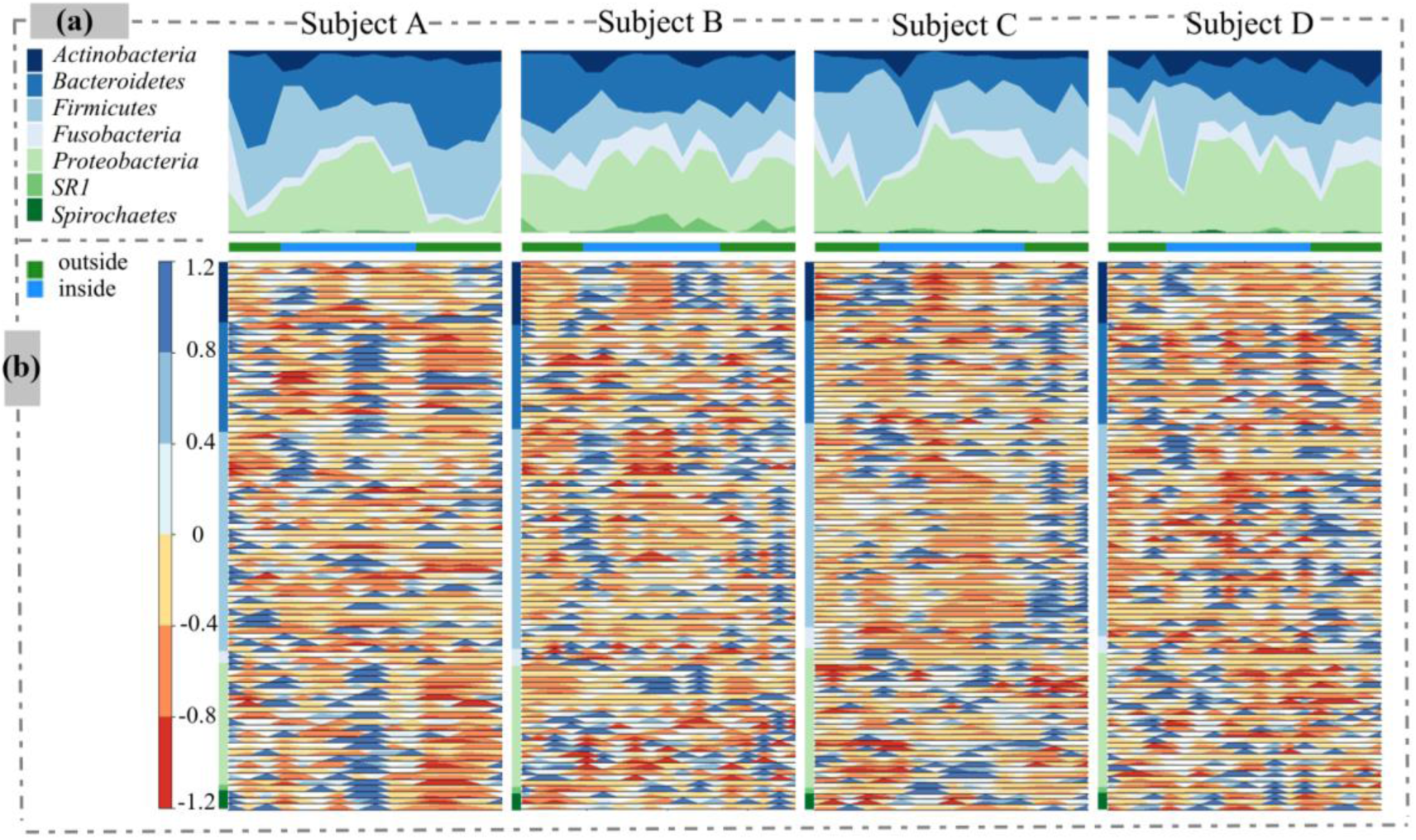
Saliva microbiota dynamics in the crewmembers throughout the experiment. (a) Percentage stacked graph showing phylum fractional abundances over time. Each ribbon represented a phylum, whose width was proportional to the phylum relative abundance at a given time point (Ribbons at the bottom of each plot indicate that the crewmembers were outside or inside the LP1). We generated the graph using R (ver 3.5.3) packages (ggplot2, plyr and reshape2). (b) Horizon graphs of the relative abundance variation of highly abundant OTUs over time. Time series were mean-centered and curves were divided into colored bands, whose width were the mean absolute deviation, that were then overlaid, with negative values mirrored upwards. Warm and cool colors indicated relative abundance below or above the mean, respectively, the darker the color, the smaller or the greater the OTU abundance. Squares on the vertical axis were colored as in (a). We generated the graph using R (ver 3.5.3) packages (latticeExtra). For the list of highly abundant OTUs, please see Supplementary Table S1.

**Fig 3:**
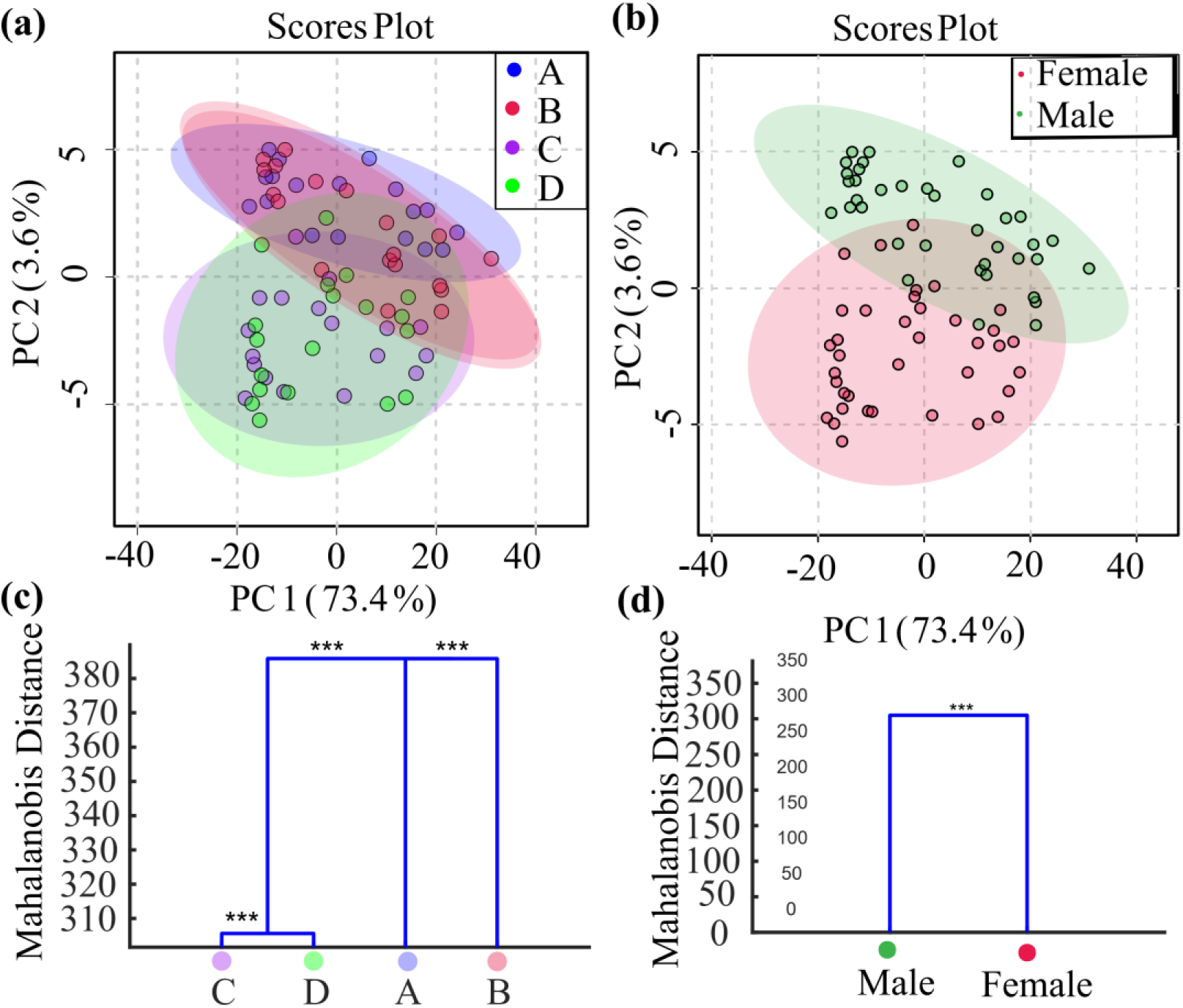
The individual and gender differences of saliva microbiome. (a-b) Principal component analyses (PCA) scores plots based on OTUs of the samples at the subject and gender, respectively. (c-d) MANOVA analysis of the different groups. The statistical significance of the separation among different groups was assessed by MANOVA test based on Mahalanobis distances using the first 25 PCs of PCA at the subject and gender. The clusters are computed by applying the single linkage method to the matrix of Mahalanobis distances between group means. *p<0.05, **p<0.01, ***p<0.001.

Besides, we adopted the autocorrelation function (ACF) to analyze the temporal dynamics of highly abundant phyla and genera in the whole experiment(McMurry and Politis 2010, 2015). Our autocorrelation analyses showed that most highly abundant phyla (Supplementary Fig S1) and genera (Supplementary Fig S2) had no significant autocorrelation, and they were stationary stochastic process, suggesting that the crewmembers’ oral microbiota had no reliable changes with time during the experiment. Furthermore, Mann-Kendall trend test(Libiseller and Grimvall 2002, McCuen 1995) was used to analyze whether there was a trending change in the continuous time period from the normal environment into LP1, the results showed that most of the trend was not significant (*P* > 0.05, Supplementary Table S2).

### Effects of controlled environment on saliva microbiota

We choose the relative abundance of the core phyla and genera to plot the curves over time, as shown in Figure 4. Results indicated that the controlled environment caused changes in the relative abundance of saliva microbiota. At phylum level, *Actinobacteria* showed a brief increase in the first week after entering LP1 (Fig 4(a)). In the first week of entering LP1, *Bacteroidetes* decreased to 19.18~31.15% of the average abundance before entering LP1, and then quickly recovered. After leaving LP1, abundance of *Bacteroidetes* returned to the previous level (except Subject B, Fig 4(a)). Besides, except for Subject D, relative abundance of *Proteobacteria* gradually increased after entering LP1 and reached 1.51~4.59 times of its relative abundance before entering LP1, then it gradually decreased and returned to the level before entering LP1(Fig 4(a)). However, change of *Firmicutes* was completely different for four crewmembers (Fig 4(a)). At genus level, *Rothia* was the most abundant of *Actinobacteria*, and its trend was consistent with *Actinobacteria* (Fig 4(b)). *Prevotella* was the most abundant of *Bacteroidetes*, and tendency of *Prevotella* was consistent with that of *Bacteroidetes*. *Streptococcus* and *Veillonella* were the most abundant of *Firmicutes*, and change of *Streptococcus* was in accordance with *Firmicutes* (Fig 4). While *Veillonella* decreased slightly after entering LP1 and recovered after leaving LP1 (Fig 4(b)). *Neisseria* and *Haemophilus* were the most abundant of *Proteobacteria*. In addition to Subject D, *Neisseria* first increased and then decreased after entering LP1, reaching the highest level about 1.73~7.28 times campared to the abundance outside of LP1 within 5 weeks (Fig 4(b)). And *Haemophilus* changed consistently with *Neisseria* but Subject B (Fig 4(b)).

**Fig 4:**
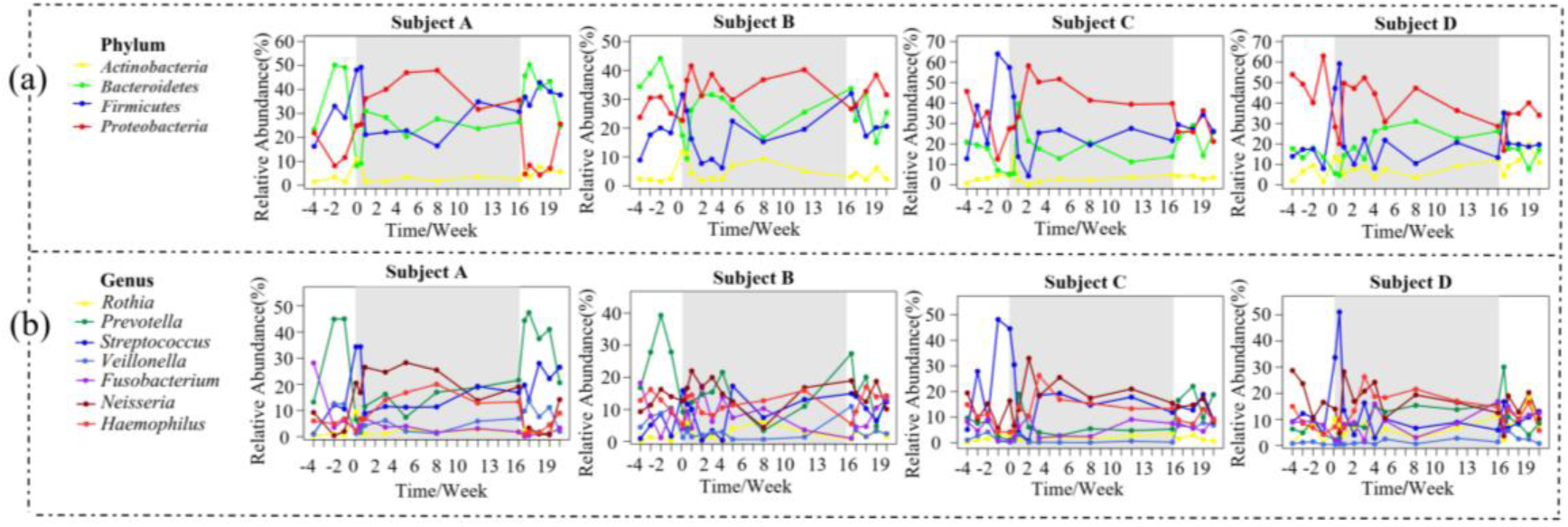
Crucial phyla and genera dynamics in crewmembers throughout the experiment. Each curve represents a phylum or genus, and shadows indicate that the crewmembers are in LP1.

### Sharing living space reduced the difference in saliva microbiota among individuals

When exploring the variation of the alpha diversity (i.e., for each crewmember) of the saliva microbiota over time, it showed apparently random fluctuations, only with large fluctuations in certain time (Supplementary Fig S3(a). And the differences among time and experimental phases were not significant (Kruskal-Wallis rank sum test, Supplementary Fig S3 (b): KW = 16.9120, P = 0.5292; Supplementary Fig S3 (c): KW = 2.3652, P = 0.3065). Furthermore, we visualized the overall changes of saliva microbiome over experimental phase using principal component analysis (PCA) (Fig 5(a)), finding that the three experimental phases could not be completely separated. We further performed MANOVA to statistically compare the differences between different experimental phases. Hierarchical clustering was generated based on mahalanobis distances as a result of MANOVA (Fig 5(b)). The MANOVA analysis showed that there were significant differences (p < 0.001) among each experimental phase. To further observe the effect of controlled environment on saliva microbiota, we compared weighted UniFrac distances among crewmembers at different time points and different phases. There was no significant difference at each time point (Kruskal-Wallis rank sum test, Fig 5(c): KW = 32.4100, P = 0.0197), but the weighted UniFrac distance decreased in the 3^rd^ Phase. It’s worthy to note that, the weighted UniFrac distances at each phase were significantly different, and the 3^rd^ Phase was significantly lower than the other two phases outside LP1 (Kruskal-Wallis rank sum test, Fig 5(d): KW = 18.1130, P = 0.0001).

**Fig 5:**
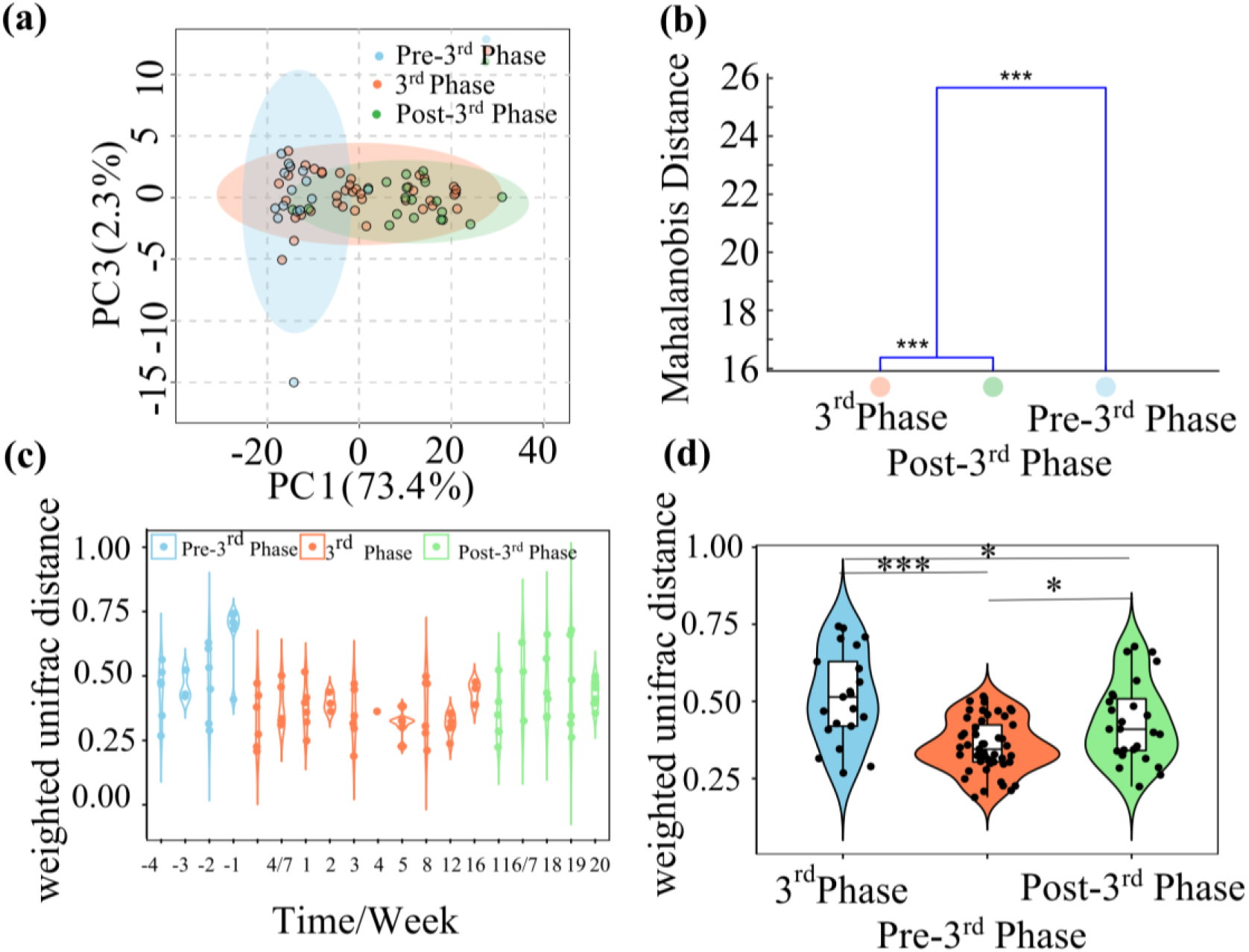
Effects of controlled environment on crewmembers’ saliva microbiota. (a) Principal component analyses (PCA) scores plots based on OTUs of the samples at the experimental phase; (b) MANOVA analysis of different experimental phases. The statistical significance of the separation among different groups was assessed by MANOVA test based on Mahalanobis distances using the first 25 PCs of PCA at the experimental phase. The clusters were computed by applying the single linkage method to the matrix of Mahalanobis distances between group means. (c-d) Violin maps of weighted UniFrac distances for crewmembers at the same time or at the same experimental phase. And differences between groups were compared by Kruskal-Wallis rank sum test. *p<0.05, **p<0.01, ***p<0.001.

### Correlation between saliva microbiota and cytokines in controlled environment

We adopted the autocorrelation function (ACF) to analyze the temporal dynamics of salivary cytokines (IL-1β, IL-6, IL-10, TNF-α and IFN-γ) in the whole experiment(McMurry and Politis 2010, 2015). Our autocorrelation analyses showed that all the cytokines had no significant autocorrelation (Supplementary Fig S4), and they were stationary stochastic process, suggesting that the crewmembers’ salivary cytokines had no reliable changes with time during the whole experiment. Furthermore, using Mann-Kendall trend test(Libiseller and Grimvall 2002, McCuen 1995) to analyze whether there was a trend change in the continuous period from the normal environment into LP1, as shown in Table 1, IL-1β showed an increasing trend (except Subject A), IL-6 (except Subject B), IL-10, TNF-α showed a decreasing trend, and IFN-γ did not change consistently. In particular, TNF-α showed a tendency to decrease significantly (P < 0.05) except Subject C.

**Table 1:** Mann-Kendall trend test of salivary cytokines. Z > 0, the time series showed an increasing trend; Z < 0, the time series showed a decreasing trend; P < 0.05, the tendency was significant.

| Cytokine | Subject | n | Mean | Z statistic | P | Sample estimates |  |  |
| --- | --- | --- | --- | --- | --- | --- | --- | --- |
|  |  |  |  |  |  | S | varS | tau |
| IL-1β | A | 22 | 5.0578 | -0.9023 | 0.3669 | -33 | 1257.6667 | -0.1429 |
|  | B | 22 | 5.8656 | 0.4512 | 0.6519 | 17 | 1257.6670 | 0.0736 |
|  | C | 22 | 4.8195 | 1.2407 | 0.2147 | 45 | 1257.6667 | 0.1948 |
|  | D | 22 | 5.6441 | 1.8047 | 0.0711 | 65 | 1257.6667 | 0.2814 |
| IL-6 | A | 22 | 10.7015 | -1.9464 | 0.0516 | -70 | 1256.6667 | -0.3037 |
|  | B | 22 | 9.6946 | 1.2407 | 0.2147 | 45 | 1257.6667 | 0.1948 |
|  | C | 22 | 9.7380 | -0.8459 | 0.3976 | -31 | 1257.6667 | -0.1342 |
|  | D | 22 | 11.8152 | -0.5076 | 0.6118 | -19 | 1257.6667 | 0.6118 |
| IL-10 | A | 22 | 3.3009 | -0.9023 | 0.3669 | -33 | 1257.6667 | -0.1429 |
|  | B | 22 | 3.1938 | -1.6919 | 0.0907 | -61 | 1257.6667 | -0.2641 |
|  | C | 22 | 3.0636 | -1.5791 | 0.1143 | -57 | 1257.6667 | -0.2468 |
|  | D | 22 | 3.4684 | -2.2558 | **0.0241** | -81 | 1257.6667 | -0.3506 |
| TNF-α | A | 22 | 17.7964 | -2.4270 | **0.0152** | -87 | 1257.6667 | -0.3783 |
|  | B | 22 | 17.5217 | -2.3686 | **0.0179** | -85 | 1257.6667 | -0.3680 |
|  | C | 22 | 16.7170 | -0.8459 | 0.3976 | -31 | 1257.6667 | -0.1342 |
|  | D | 22 | 17.9889 | -2.9890 | **0.0028** | -107 | 1257.6667 | -0.4632 |
| IFN-γ | A | 22 | 2.2153 | -1.5227 | 0.1278 | -55 | 1257.6667 | -0.2381 |
|  | B | 22 | 2.1919 | 0.0000 | 1.0000 | -1 | 1257.6667 | -0.0043 |
|  | C | 22 | 1.8623 | -2.1721 | **0.0299** | -78 | 1257.6667 | -0.3384 |
|  | D | 22 | 2.3966 | 0.5924 | 0.5536 | 22 | 1257.6667 | 0.0954 |

We analyzed the Spearman correlation between highly abundant salivary microbial genera and salivary cytokines of each crewmember in LP1 (Supplementary Table S3), as shown in Fig 6. Co-occurrence network of four crewmembers have strong individual characteristics, which of Subject C was the most complex and showed more complicated correlation among saliva microbiota and cytokines. It was worth noting that, in addition to Subject D, TNF-α showed a consistent positive correlation with *Actinomyces* and *Rothia* (R ≥ 0.5).

**Fig 6:**
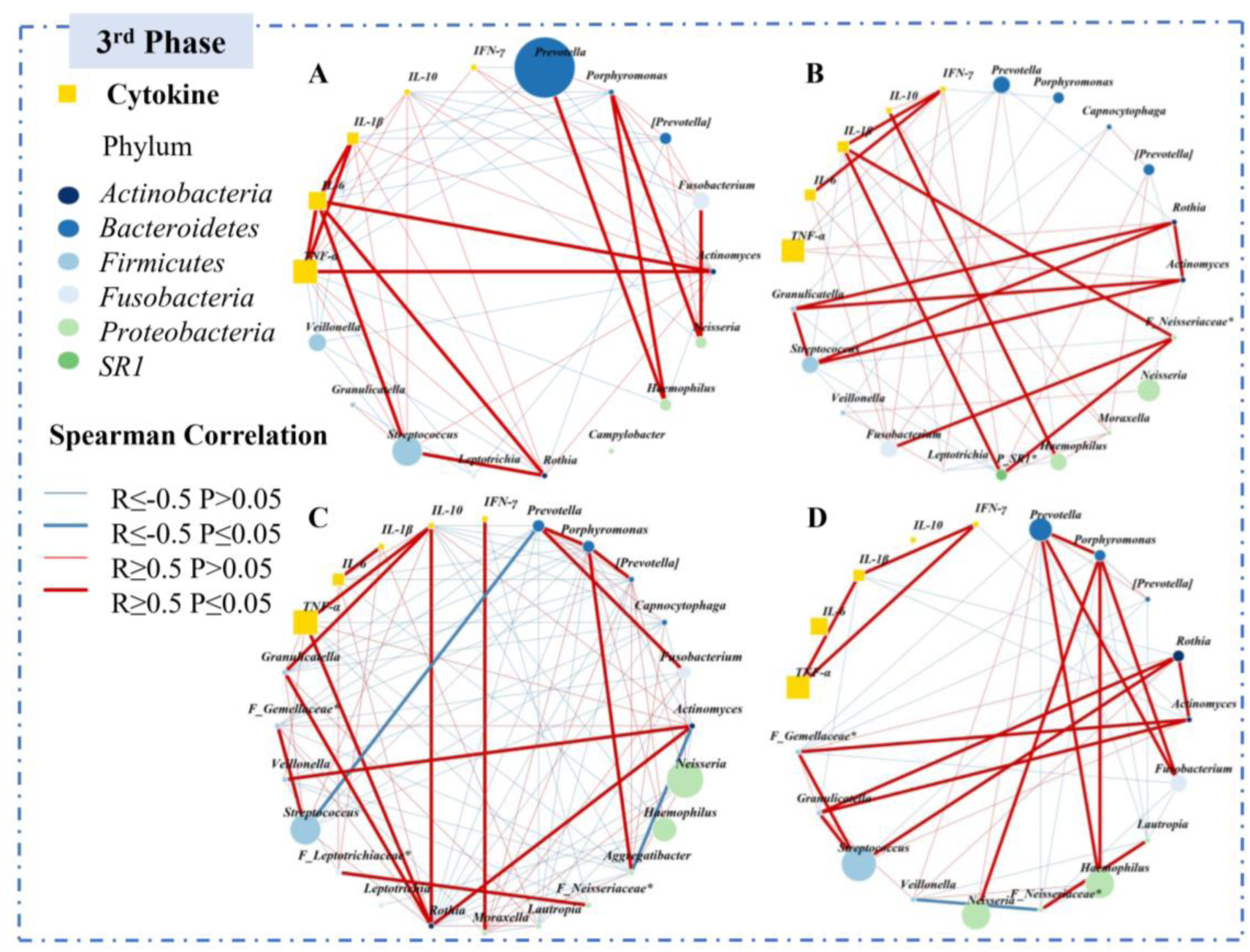
Co-occurrence network of crewmembers’ salivary cytokines and highly abundant microbial genera in LP1: Select genus with relative abundance over 1%, calculate the spearman correlation coefficient of saliva microbiota and cytokines in LP1, and then generate the graph using R (ver 3.5.3) packages (igraph). Each node represented a microbial genus or a salivary cytokine. Circles colored by phylum represented different genus, and squares represented different cytokines. The size of the nodes was proportional to the microbial abundance or salivary cytokine level. The link between nodes indicated spearman correlation (|r|≥0.5). The blue and red lines indicate negative correlation and positive correlation, respectively. Particularly, bold indicated significant correlation (p≤0.05).

## Discussion

During long-term space missions, health problems such as symbiotic microbial disorders and impaired immunity often occur because of psychological and environmental pressures, which in turn affects space missions. The ‘‘Lunar Palace 365” project was a multicrew and closed experiment carried out in LP1, representing an invaluable opportunity to study symbiotic microbial disorders and immune adaptation problems that humans may face in long-term space missions.

Within this context, we explored the temporal dynamics of the saliva microbiota and salivary cytokines in the four crewmembers participating in the 3^rd^ Phase of “Lunar Palace 365”, across the 3^rd^ Phase of the project, including the period before entering LP1, and after the return to regular life, and about 6 months of sampling. Previous studies have found changes in oral microbiota(Brown et al. 1976), infectious diseases(Taylor and Sommer 2005) and decreased immunity(Mermel 2013) in space missions. In the past, researchers thought that was mainly caused by cosmic radiation, microgravity and long-term consumption of pre-packaged food. In the “Lunar Palace 365” project, we kept the crewmembers’ diet consistent with regular life by planting food and vegetables, and studied the effects of closed semi-sterile environment on human saliva microbiota and oral immunity for the first time.

Saliva microbiota is more stable than other symbiotic microorganisms(Costello et al. 2009) and dosen’t change significantly within 1 year for adult(Cameron et al. 2015). Our results are consistent with previous studies, and the dominant species steadily fluctuate during the experiment, with no significant trendency (Supplementary Fig S1, Fig S2 and Table S1). Besides, there was no significant difference in the alpha diversity of the four crewmembers at different time and different phases (Supplementary Fig S3), indicating that the crewmembers were well adapted to the closed environment and the salivary microbial community remained stable. Though the oral microbiome of single person is relatively stable, it’s significantly different among individuals(Jakubovics 2015). Nasidze et al.(Nasidze et al. 2009) analyzed saliva samples of 120 healthy people from 12 countries and regions worldwide and found the oral microbiota was significantly individual specific. Our results are consistent with previous studies, and the oral microbial communities of the four crewmembers show significant individual and gender differences.

However, we also found some microorganisms showed transient trends in the closed environment, including the reduction of *Bacteroidetes* and the increase of *Proteobacteria* (Figure 4 (a)). A study based on 16S pyrosequencing found that oral *Bacteroidetes* was significantly elevated in patients with periodontitis, while *Proteobacteria* was reduced(Griffen et al. 2012). In addition, Said et al.(Said et al. 2013) found that patients with inflammatory bowel disease had more abundant salivary *Bacteroidetes*, accompanied with reduced *Proteobacteria*. Based on previous research, we believe that trends of highly abundant phyla seem to be beneficial to oral and general health in the closed environment. Moreover, we found changes in phylum were mainly caused by highly abundant genus. The changes of *Rothia*, *Prevotella* and *Streptococcus* were consistent with *Actinobacteria*, *Bacteroidetes* and *Firmicutes*, respectively. And trends of *Neisseria* and *Haemophilus* were roughly the same as that of *Proteobacteria*. *Prevotella* existed extensively in human microbiota, and significantly increased in saliva of patients with caries(Yang et al. 2009), esophagitis(Yang et al. 2009), sinusitis(Li et al. 2009), inflammatory bowel disease(Said et al. 2013), HIV with hyperviralemia(Dang et al. 2012) and bacterial vaginosis(Oakley et al. 2008). In our results, *Prevotella* was reduced in controlled environment except for Subject D, indicating well health of crewmembers in controlled environment. In addition, we found *Neisseria* and *Haemophilus* were elevated in controlled environment, which were associated with type 2 diabetes(Casarin et al. 2013) and oral leukoplakia(Hu et al. 2016), respectively. We suspect that it may be related to simpler food processing in the controlled environment. Several studies discovered that oral microbes of hunting people, traditional farmers, Westerners and vegetarians varied greatly, and finely processed foods resulted in less abundent *Neisseria* and *Haemophilus*(Clemente et al. 2015, Lassalle et al. 2018). Dietary changes in long-term mission play an important role in saliva microbiota. The increased *Streptococcus* in astronauts’ saliva during Skylab missions was thought to be caused by intakeing of prepared foods (Brown et al. 1976). However, our study did not show similar results, indicating that ensuring the supply of fresh food by BLSS helped reduce adverse changes in oral microbiota.

In addition, according to the weighted UniFrac beta diversity, the bacterial communities of the four crewmembers became, to some extent, more similar to each other over time, suggesting a certain degree of convergence of the temporal dynamics of abundant microbiota taxa in humans sharing a confined environment. Our previous 105-day closed experiment(Hao et al. 2018) and the Mars 500 ground-based space simulationt(Turroni et al. 2017) found similar phenomena in human intestinal microbes. Our result is similar to previous studies, although saliva microbiota has strong individual characteristics, small-scale effects due to shared living spaces can significantly affect the composition of microbial communities(Shaw et al. 2017). Stahringer et al.(Stahringer et al. 2012) observed the same effect in the salivary microbiome and also found that the salivary microbiomes of twins became less similar as they grew older and ceased cohabiting, concluding that “nurture trumps nature” in the salivary microbiome. Our work supporting the dominant role of the environment in affecting salivary microbiome composition suggests that another important factor in long-term persistence may be the regular reseeding of the ecosystem with bacteria from the external environment. This also reminds us to pay attention to crewmembers saliva microbiota and environmental microbes in long-term space missions. And it is necessary to prevent the breeding of harmful microorganisms in the space.

Cytokines are important components in saliva, participating in host defenses to maintain oral and systemic health, and providing information on local and system conditions(Farnaud et al. 2010). Our study found that salivary cytokines fluctuate smoothly throughout the experiment. However, when the crewmembers entered the controlled environment from normal environment, IL-1β showed an increasing trend, IL-6 and IL-10 showed a decreasing trend, and TNF-α showed a significant downward trend (Table 1). Changes in salivary cytokines may be caused by oral microbes. Increased periodontal pathogens will enhance stimulations to TLRs, induce Th1 to secrete IL-2, IFN-γ, TNF-α to kill intracellular infection pathogens, and then induce Th2 secretion of IL-4, IL-5, IL-6, IL-13 to regulate humoral immunity and limit Th1 response(Lazarevic et al. 2010). On the other hand, oral symbiotic microorganisms can induce the secretion of IL-10 by T-reg cells and inhibit the activity of effector T cells(Loesche et al. 1975). The crewmembers’ salivary cytokines showed a decreasing trend in the closed environment, indicating there was no pathogenic change in saliva microbiota and didn’t cause abnormal inflammatory response.

In addition, IL-1β is a pro-inflammatory cytokine stimulated by bacterial lipopolysaccharide, which participates in the inflammatory response and stimulates immune cells to secrete IL-1β, IL-2, IL-6, IL-8, TNF-α and IFN-γ(Kampoli et al. 2009). In our study, elevated IL-1β may be caused by psychosocial stress and increased negative emotions. Schbacher et al.(Aschbacher et al. 2009) found the response of IL-1β to stress had predictive validity for mental health. Szabo et al. (Szabo et al. 2019) also found that psychosocial stress was associated with higher salivary IL-1β, and increased negative emotions caused an increase in IL-1β(Newton et al. 2017). Furthermore, maintaining positive emotions can alleviate negative effects of stress by reducing the inflammatory response (Fredrickson 2004). Although the crewmembers seemed not to experience psychological distress in the controlled environment(Hao et al. 2019), Mars 500 has reported symptoms of depression in 93% of mission weeks in a 520-day ground space simulation experiment(Basner et al. 2013, Basner et al. 2014), and found serum serotonin significantly elevated in the mid-late stage of the mission(Wang et al. 2014). Serotonin plays an important role in regulating stress(Christine and C Rob 2007) and enhancing psycho-physiological resistance to chronic stress(Grippo et al. 2005, Porter et al. 2004, Storey et al. 2006). Driven by the defensive system, unpleasant emotions, such as anxiety, depression and fear, induce different reflexive automatic and somatic outputs(Koganemaru et al. 2012, Lang and Bradley 2010, Smith et al. 2005). In the current study, when the psychological status of the crewmembers became worse with time, they expressed withdrawal from the negative stimuli with a positive rating bias. As a result, the secretion of crewmembers’ salivary IL-1β increased from normal environment to controlled environment, while the subjective evaluation of the crewmembers showed a good emotional state in the whole experiment.

Strong correlations between some inflammatory biomarkers and saliva microbiota compositions has been revealed, for instance, IL-17 plays an important role in oral antifungal(Wade 2011). Besides, Said et al.(Said et al. 2013) found lower lysozyme and elevated IL-1β, IL-8, IgA in saliva of patients with inflammatory bowel disease were likely to be synergistically or interactively associated with the abundance of *Streptococcus*, *Prevotella*, *Veillonella*, and *Haemophilus*. *Porphyromonas gingivalis* can be colonized in atherosclerotic plaques, stimulating the secretion of inflammatory factors such as TNF-α, IL-1β, IL-6, and causing inflammation and vascular endothelial damage(Fåk et al. 2015). We found complex correlations among salivary cytokines and highly abundant genera in the controlled environment, and TNF-α showed a consistent correlation with *Actinomyces* and *Rothia*. *Actinomyces* is an early colony in the process of plaque maturation. Sato et al.(Sato et al. 2012) found peptidoglycans of *Actinomyces naeslundii* can cause the production of IL-1β, IL-6 and TNF-α. In our study, the positive correlation between TNF-α and *Actinomyces* confirms their research, and also suggests that decreased TNF-α in saliva of crewmembers in a controlled environment may be related to decreased *Actinomyces*. Beaides, *Rothia* is a conditional pathogen widely present in human oral cavity, and the most typical species is *Rothia dentocariosa*. Kataoka et al.(Kataoka et al. 2013) found that *R. dentocariosa* can induce TNF-α production through TLR2, which is consistent with our findings. Our study suggests that there is also certain correlation between saliva microbiota and salivary cytokines in healthy individuals, but disturbed by the daily life environment. When studying the correlation between saliva microbiota and salivary cytokines, we should consider hosts’ health status, diet and living environment to make the results more accurate.

In conclusion, our study proves that the closed isolation semi-sterile environment does not cause systemic microorganisms and immune disorders, and saliva microbiota and salivary cytokines fluctuate smoothly throughout the experiment. Although salivary IL-1β has a tendency to increase in the controlled environment, it is more likely to be a stress response under prolonged stress. Overall, crewmembers’ salivary inflammatory cytokines are reduced in the controlled environment. In the whole experiment, although salivary microorganisms and salivary cytokines transiently fluctuated after entering the controlled environment, most of them returned to previous levels after back to the normal environment, indicating that crewmembers adapted well to the closed isolation environment without permanent changes. Besides, it provides a new idea for future research on the impact of closed isolation cabins on crewmembers health, and provides guidance for studying the effect of semi-sterile environments on human immunity based on saliva microbiota.

## Compliance with ethical standards

### Ethics approval and consent to participate

We obtained written informed consent from four subjects enrolled in the study. This study was approved by the Science and Ethics Committee of School of Biological Science and Medical Engineering in Beihang University,Beijing, China (Approval ID: BM20180003) and complied with the Helsinki Declaration.

### Author contributions

Y.Z., C.D. and H.L. designed the study. H.L. supervised the study. Y.Z. performed the experiments. Y.Z., Z.H, Y.F., and J.Y. performed data analysis. Y.Z. and Z.H. wrote the paper. Y.Z., Z.H, Y.F., J.Y., C.D. and H.L. contributed to the editing and revision of the paper. All authors read and approved the final manuscript.

### Associated Content

Supplementary materials (PDF, 967kb)

Supplementary materials (Excel, 37kb)

